# Morphologic Classification and Automatic Diagnosis of Bacterial Vaginosis by Deep Neural Networks

**DOI:** 10.1101/2020.05.20.101055

**Authors:** Zhongxiao Wang, Lei Zhang, Min Zhao, Ying Wang, Huihui Bai, Yufeng Wang, Can Rui, Chong Fan, Jiao Li, Na Li, Xinhuan Liu, Zitao Wang, Yanyan Si, Andrea Feng, Mingxuan Li, Qiongqiong Zhang, Zhe Yang, Mengdi Wang, Wei Wu, Yang Cao, Lin Qi, Xin Zeng, Li Geng, Ruifang An, Ping Li, Zhaohui Liu, Qiao Qiao, Weipei Zhu, Weike Mo, Qinping Liao, Wei Xu

**Affiliations:** Department of Obstetrics and Gynecology, Beijing Tsinghua Changgung Hospital, School of Clinical Medicine, Tsinghua University, Beijing 102218, China; Institute for Interdisciplinary Information Sciences, Tsinghua University, Beijing 100084, China; Suzhou Turing Microbial Technologies Co., Ltd, Suzhou 215021, China; Beijing Turing Microbial Technologies Co., Ltd, Beijing 101318, China; The Second Affiliated Hospital of Soochow University, Suzhou 215004, China; The Affiliated Hospital of Inner Mongolia Medical University, Hohhot 010050, China; Beijing Obstetrics and Gynecology Hospital,Capital Medical University Beijing Maternal and Child Health Care Hospital, Beijing 100000, China; Peking University First Hospital, Beijing 100035, China; Women’s Hospital of Nanjing Medical University, Nanjing Maternity and Child Health Care Hospital, Nanjing 210004, China; The First Affiliated Hospital of Xi’an Jiaotong University, Xi’an 710061, China; Peking University Third Hospital, Beijing 100191, China; Department of Operations Research and Financial Engineering, Princeton University, Princeton, NJ 08544; Beijing HarMoniCare Women’s and Children’s Hospital, Beijing 100029, China; Binzhou Medical University Hospital, Binzhou 256600, China; Department of Physics, Tsinghua University, Beijing 100084, China; School of Clinical Medicine, Tsinghua University, Beijing 100084, China

**Author notes:** These two authors contributed equally (Zhongxiao Wang, Lei Zhang).

**Keywords:** Application of AI to diagnostic microbiology, Automation in clinical microbiology

## Abstract

**Background:** Bacterial vaginosis (BV) was the most common condition for women’s health caused by the disruption of normal vaginal flora and an overgrowth of certain disease-causing bacteria, affecting 30-50% of women at some time in their lives. Gram stain followed by Nugent scoring (NS) based on bacterial morphotypes under the microscope was long considered golden standard for BV diagnosis. This conventional manual method was often considered labor intensive, time consuming, and variable results from person to person.

**Methods:** We developed four convolutional neural networks (CNN) models, and evaluated their ability to automatic identify vaginal bacteria and classify Nugent scores from microscope images. All the CNN models were first trained with 23280 microscopic images labeled with Nugent scores from top experts. A separate set of 5815 images were evaluated by the CNN models. The best CNN model was selected to generalize its application on an independent sets of 1082 images collecting from three teaching hospitals. Different hardwares were used to take images in hospitals.

**Results:** Our model could classify three Nugent Scores from images with high three classification accuracy of 89.3% (with 82.4% sensitivity and 96.6% specificity) on the 5815 test images, which was better diagnostic yield than the top-level technologists and obstetricians in China. The ability of generalization for our model was strong that it obtained 75.1%, which was 6.6% higher than the average of technologists.

**Conclusion:** The CNN model over performed human healthcare practitioners on accuracy, efficiency and stability for BV diagnosis using microscopic image-based Nugent scores. The deep learning model may offer translational application in automating diagnosis of bacterial vaginosis with proper supporting hardware.

## Introduction

Abnormal vaginal discharge and odor were vaginitis symptoms that affected millions of women globally and represented the most commen reasons for women to visit clinics. Bacterial vaginosis (BV, 40-50%), vulvovaginal candidiasis (VVC, 20-25%) and trichomoniasis (TV, 1.5-2.0%) were the leading causes of vaginitis. Bacterial vaginosis (BV) represented a dysbiosis of the vaginal microbiome that was associated with significant adverse healthcare outcomes, including preterm labor resulting in low birth weight, pelvic inflammatory disease, acquisition of the human immunodeficiency virus and increased susceptibility to sexually transmitted infections [1–8]. In the United States, women had a high BV incidence rate of 29.2% with the prevalence varies with race: African-American (51%), Hispanic (32%), and Whites (23%) [9]. In China, the prevelance of BV in a few cities with survey data was between 15-20%, which represented more than 100 million women [10].

Different methods were developed for diagnosing BV, but gram-stained microscopy was considered gold standard method [11]. Clinical criteria for diagnosis of BV only had a sensitivity of 60-72% [12,13]. Other methods including enzymatic test such as the OSOM BV Blue, and molecular method such as BD MAX™ Vaginal Panel had higher sensitivities of 91.7% and 90.7% respectively for the diagnosis of BV [14,15]. Therefore, The Guidelines Group recommended that the current best test to diagnose BV in women was gram-stained microscopy [11].

In 1991, Nugent et al [16] reported the use of a numerical score to diagnose BV by semiquantization of gram-positive rods, gram-negative coccobacilli forms, and curved gram-negative rods after Gram staining. These morphotypes were thought to represent Lactobacillus spp., Gardnerella vaginalis and Mobiluncus spp., respectively. Nugent scoring had since then become the ‘gold standard’ for laboratory diagnosis of BV [7, 8, 17]. In Nugent scale, scores of 0–3 were considered to have normal vaginal flora (Lactobacillus dominant); scores of 4–6 were labeled as altered vaginal flora (mixed morphotypes); and scores of 7–10 were indicative of BV (absence of lactobacilli and predominance of the other 2 morphotypes). Alternative diagnostic methods such as molecular diagnostic assays, enzymatic assays, and chromogenic point-of-care test (POCT) were compared to Nugent criteria [18]. However, the determination of a Nugent score by a microbiologist could be easily influenced by individual skill and was time consuming [17, 18]. In addition, the number of experienced microbiologists or technologist performing the microscopic work was in shortage among different countries and districts [19]. Needed was a more efficient method to classify Nugent scores.

Here, we provided a proof of concept for a deep-learning-based model to quantify Gram stain and, hence, automated classification of Nugent scores. Recently, a traditional image processing method was developed for automatic bacterial vaginosis diagnosis. However, the sensitivity (58.3%) and specificity (79.1%) was relatively poor in comparison to expert due to its limitation in the image processing algorithm [17]. Deep learning method especially convolutional neural network (CNN) models had demonstrated excellent performance on computer vision tasks including image classification, image semantic segmentation and image object detection. For the image classification, various CNN models were constructed with increasing performance on natural image classification. Numerous CNN models, including LeNet-5 [20], AlexNet [21], VGGNet [22], ResNet [23], GooLeNet [24], Xception [25], FCN [26], PSPNet [27], and U-net [28], were developed to improve performance on natural image recognition. Many models were proved effective for medical image processing, from identifying diabetic retinopathy in retinal fundus photographs [29–31], endoscopic images [32], to microbiology recognitions [33, 34]. We hypothesized that CNN based deep learning models can be used to diagnose BV using Nugent score classifications efficiently and accurately. First we developed several CNN models to learn images previously diagnosed and curated by obestetricians and microbiologiests. Second, a trained model were used to test on a separate image set from the same hospital. Our trained model was subsequently evaluated for accuracy in comparison to expert classification. Finally, three independent test sets collected from three different medical institutions were used to verify the versality of our CNN model.

## 2 Material and Methods

### 2.1 Image Data Preparation

A total of 29095 microscopic images including associated medical records from January, 2018 to September, 2019 at Beijing Tsinghua Changgung Hospital were retrieved. One-fifth of all samples were randomly selected as the test set (5815 samples), and the rest samples (23280 samples) were used as the training set. The resolution of the samples was 1024 × 768 pixels.

For verifying our model’s accuracy in extensive setting and comparing it with various experts, three independent testing data sets Γ, Λ, Σ were constructed. 427 images randomly selected from the original 5815 test sets were Set Γ; 359 images collected from The Second Affiliated Hospital of Soochow University were Set Λ; and 296 samples from The Affiliated Hospital of Inner Mongolia Medical University were Set Σ. The resolution of the samples in sets Λ and Σ was 1280 × 1024 pixels. To standardize the images, the center 1280 × 960 pixels were cropped and resized to 1024 × 768 pixels.

The diagnosis of all images was made by the experts from the National Committee of Gynecological Infection, Chinese Medical Association including two chief obestetricians and three microbiologists based on microscopic images. Each sample was firstly diagnosed by two microbiologists, with discordant samples then reviewed by the chief obstetrician to make final decision. The labeled Nugent scores ranged from 0 to 10, which were divided into three groups: normal vaginal flora (0-3 scores), altered vaginal flora (4-6 scores) and BV(7-10 scores).

Three representative images and their distributions of Nugent classifications from the three different medical institutions above were shown in Fig.1. The training set of 23280 images contained 16490 images with normal vaginal flora, 4660 images with altered vaginal flora, and 2130 images diagnosed with BV. The test set of 5815 samples included 4120 images scored 0-3, 1164 images scored 4-6, and 531 images scored 7-10. Set Γ had 160 normal vaginal flora, 206 altered vaginal flora and 61 BV images. Set Λ had 109 normal vaginal flora, 149 altered vaginal flora and 101 BV images. Set Σ had 158 normal vaginal flora, 123 altered vaginal flora and 15 BV images. The distributions of Nugent classifications for each set were shown in Tab1.

**Table 1:** The distributions of Nugent classifications for each set.

| Data Sets | Normal<br>Vaginal<br>Flora (0-3) | Altered<br>Vaginal<br>Flora (4-6) | BV (7-10) | Total |
| --- | --- | --- | --- | --- |
| Training Set | 16490 | 4660 | 2130 | 23280 |
| Test Set | 4120 | 1164 | 531 | 5815 |
| Test Set $\Gamma$ | 160 | 206 | 61 | 427 |
| Test Set $\Lambda$ | 109 | 149 | 101 | 359 |
| Test Set $\Sigma$ | 158 | 123 | 15 | 296 |

**Figure 1:**
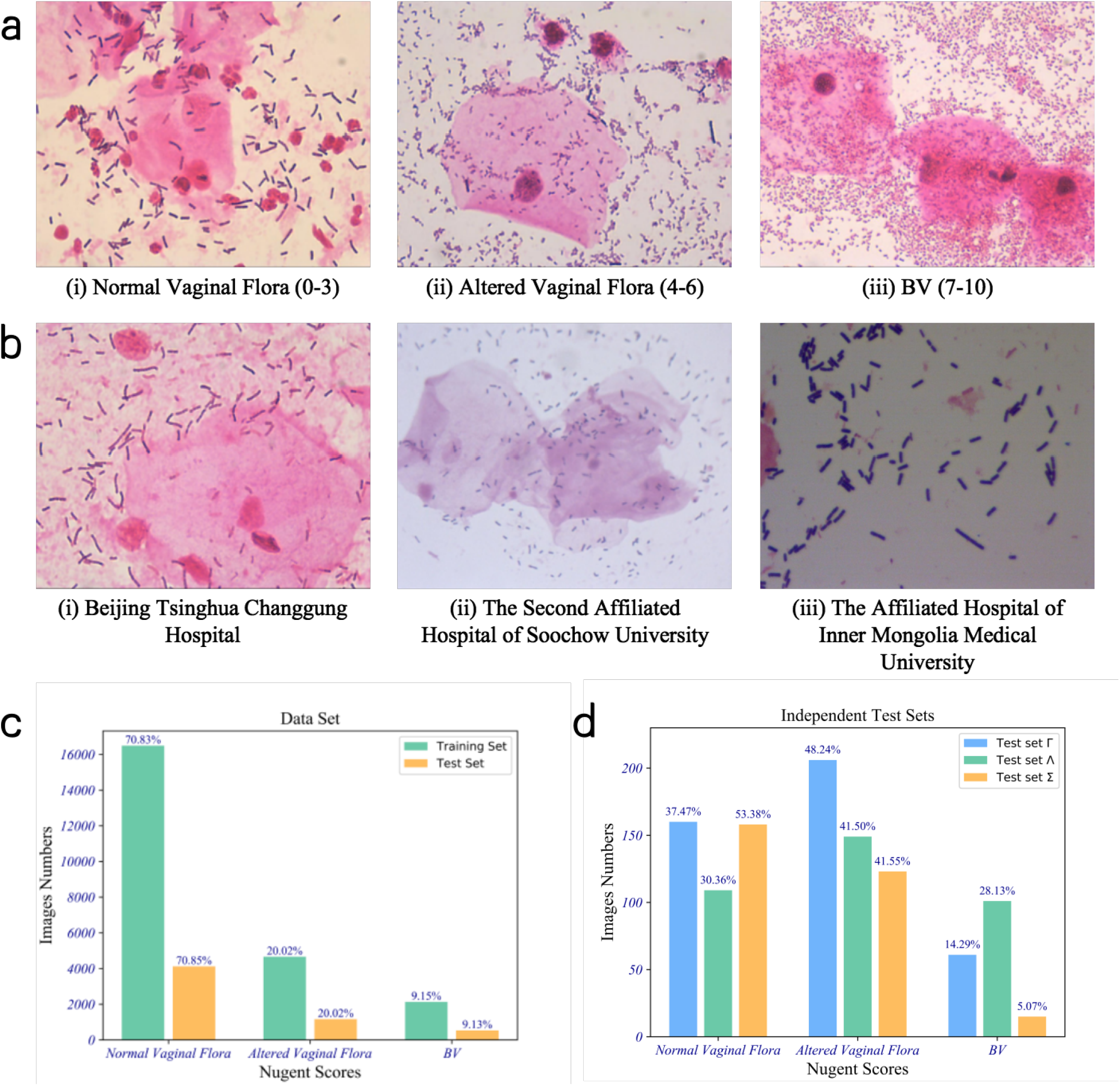
The data set information. a) Three typical samples for (i) Normal Vaginal Flora, (ii) Altered Vaginal Flora, and (iii) BV collecting from Beijing Tsinghua Changgung Hospital. b) Three typical samples collecting from (i) Beijing Tsinghua Changgung Hospital, (ii) The Second Affiliated Hospital of Soochow University, and (iii) The Affiliated Hospital of Inner Mongolia Medical University. c) The distribution of the data set. d) The distribution of the three independent test sets Γ from Beijing Tsinghua Changgung Hospital, Λ from The Second Affiliated Hospital of Soochow University, and Σ from The Affiliated Hospital of Inner Mongolia Medical University.

The study protocol was approved by the Ethics Committees of participating hospitals.

### 2.2 Development of CNN Models for BV Analysis

We developed four CNN models with different network widths (different channel numbers in each layers) to predict Nugent scores based on microscope images. The residual module used in ResNet was employed in all models [23]. In the training process, color jittering, scale jittering, horizontal flip and vertical flip were used as the data augmentation methods. By comparing the performance of the four models, the best model with the best network width was selected.

In clinical practice, the microbiologist/technologist inspected multiple fields of view for each sample under the microscope. Each field of view was designated a Nugent score. A final diagnostic result was based on collective Nugent scores from various fields. A representative image of the diagnostic result would be selected and saved. Our model was applied to the representative images for an automated Nugent score to achieve automatic diagnosis.

The classic classification convolutional neural network was unsuitable to process our microscope images at 1024 × 768 resolution, as they are for input pictures with a resolution of 224 × 224, such as VGG, GooLeNet, ResNet and DenseNet [21-24]. If we directly resized the images to 224 × 224, lots of useful information would be lost or altered, especially the shapes and details of many bacteria used for Nugent score. Therefore, we developed a new CNN model named NugentNet to adapt the input of our microscope images (Fig.2).

**Figure 2:**
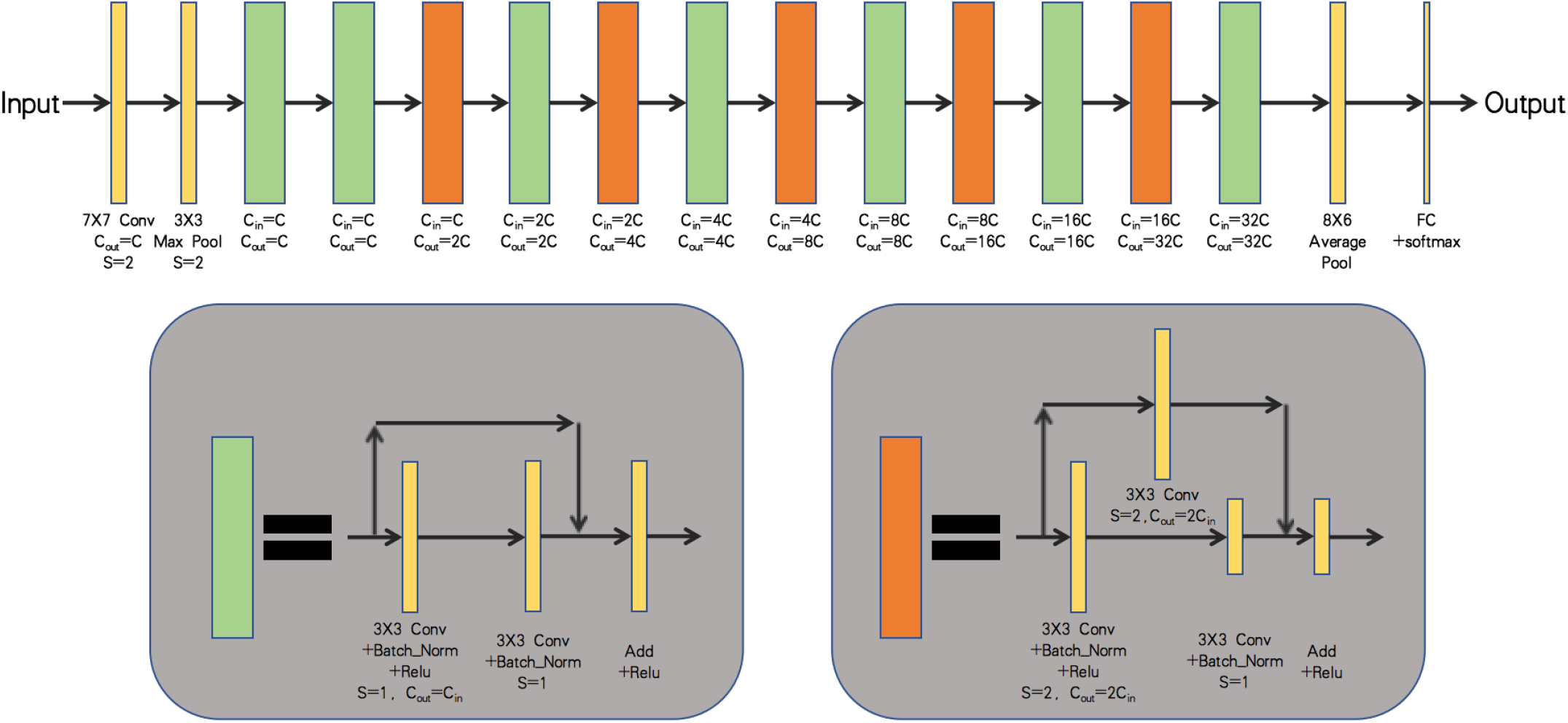
The architecture of our models. Conv represented convolutional layer and batch norm was the batch normalization layer. S represented the stride of the convolution operation. FC was the fully connected layer. *C_in_* and *C_out_* were the input channel and output channel. *C* determined the width of the network. For the basic model *C* = 64.

The model was trained to minimize a cross entropy loss function given by

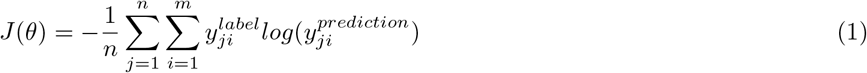

where *m, n* were the number of the classes and the batch size; *y^label^* was the one-hot encode vector of the label; 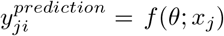 was a vector with the elements that represented the probabilities for predicting each class, which was obtained by using a softmax after the last fully connected layer in the CNN model. *x_j_* was the input data and *θ* were the variables for updating. Momentum optimizer was used [35] to train the model on the labeled images.

The training set only had 23280 samples. But the basic model needed more samples to train and it showed overfit on the data set. Therefore, we further developed three compression model, needed less training samples than the basic model, to find the most suitable model for our data set.

The compression models (1/2 NugentNet, 1/4 NugentNet, 1/8 NugentNet) were derived from the basis model. The value of C in Fig.2 were reduced to 32 for 1/2 NugentNet, 16 for 1/4 NugentNet and 8 for 1/8 NugentNet, which means the input and output channels for every convolutional layers were reduced to 1/2, 1/4 and 1/8 of the basic model. Reduced channels would simply the neural network and hence reduced the use of computing resources. Therefore, the speed of compression models was quicker than the basic model.

### 2.3 Image Preprocessing for CNN Models

The three independent test sets Γ, Λ, Σ collected from different hardwares used by the above three hospitals to generate microscope images (Tab.2). Three different typical samples were collected from the three hospitals (Fig.1(b)): Set Γ from Beijing Tsinghua Changgung Hospital (Fig.1(b[i]), Set Λ from The Second Affiliated Hospital of Soochow University (Fig.1(b[ii]), and Set Σ collected from The Affiliated Hospital of Inner Mongolia Medical University (Fig.1(b[iii]). The pixel distributions of these three types of samples were significantly different because they were in different settings, by different people, and from different hardwares. The main differences among three test sets were the actual physical area represented by the image and the brightness of the image (Tab.2). The actual physical area of images in Λ was twice that of the sample in the training set. Images in Set Λ were also brighter than images in the training set. In contrast, the images in Σ represented only half actual physical area and darker than the training set.

**Table 2:** Different hardwares used by different hospitals to generate microscope images and the main differences of the samples.

| Hospital | Camera Models | Camera Sensor Size | C-Mount Magnification | Target Physical Area | Resolution | Test Set Pixel Average Values (B,G,R) |
| --- | --- | --- | --- | --- | --- | --- |
| Beijing Tsinghua Changgung Hospital | MC170HD | $6.16mm \times 4.62mm$ | $0.7\times$ | $87.02um \times 65.26um$ | $1024 \times 768$ | (187, 172, 226) |
| The Affiliated Hospital of Inner Mongolia Medical University | UI-3240LE-C-HQ | $7.18mm \times 5.32mm$ | $1\times$ | $67.84um \times 54.27um$ | $1280 \times 1024$ | (155, 111, 117) |
| The Second Affiliated Hospital of Soochow University | UI-3240LE-C-HQ | $7.18mm \times 5.32mm$ | $0.5\times$ | $135.68um \times 108.54um$ | $1280 \times 1024$ | (208, 181, 194) |

Preprocessing was used to eliminate sample differences among different test sets in the inference process. The preprocessing included two steps: standardizing the actual physical area of images, and adjustment of the brightness of images (Fig.3). The preprocessing for test set Λ was shown in Fig.3(a). First, the center 656 × 492 pixels were cropped and resized to 1024 × 768 pixels. Second, all pixel values were increased by 32 for red channel, decreased 9 for green channel and decreased 21 for blue channel. Shown in Fig.3(b), the preprocessing of test set Σ started by resizing to 798 × 598 pixels, then followed by edge expansion. Three typical edge expansion methods: replicate, wrap and reflect were applied (Fig.3(c)). Our results showed the reflect method was the best. All pixel values of images were increased 109 for red channel, 61 for green channel and 32 for blue channel in Set Σ.

**Figure 3:**
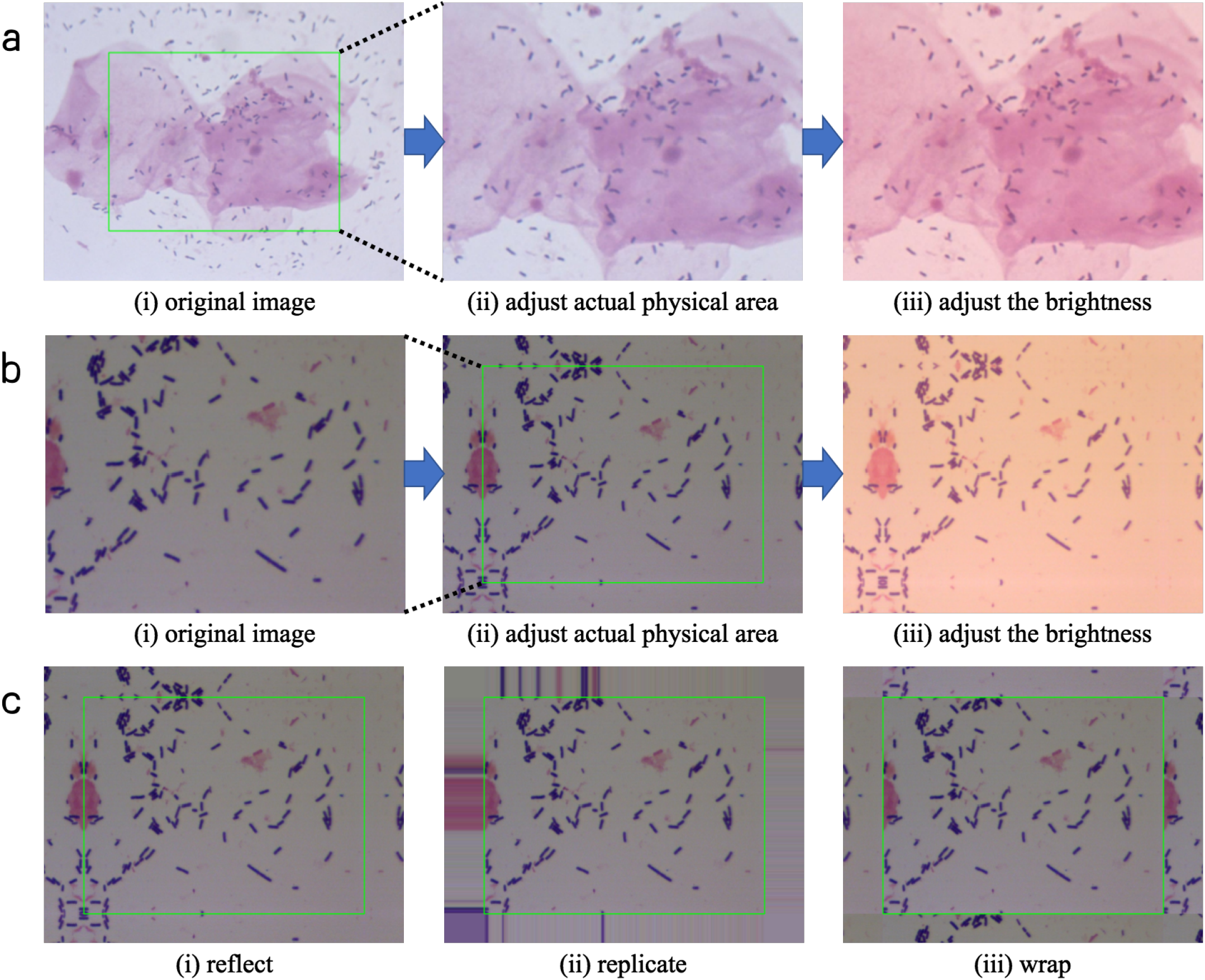
The preprocessing for test set Λ and Σ. a) was the preprocessing for Λ. b) was the preprocessing for Σ. c) was three typical edge expansion methods used in b[ii].

### 2.4 Analysis of Diagnostic Performance Using Metrics Methods

The performances of diagnostic methods were usually measured by sensitivity and specificity in comparison to the standard method. The sensitivity represents the true positive rate, while the specificity represents the true negative rate. The two diagnostic indexes was calculated by

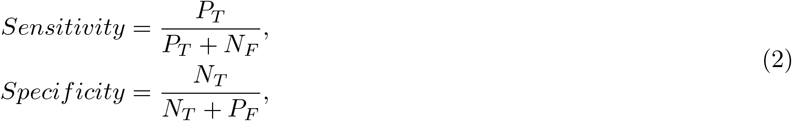

where *P_T_* was the number of the positive samples being correctly diagnosed (True Positive), *N_F_* was the number of the positive samples being diagnosed as negative (False Negative), *N_T_* was the number of the negative samples being correctly diagnosed (True Negative), *P_F_* was the number of the negative samples being diagnosed as positive (False Positive). In this study, we considered normal vaginal flora as the negative samples, altered vaginal flora and BV as the positive samples.

The performance of our models was illustrated by AUC (area under receiver operating characteristic (ROC) curve). ROC curve was a graphical plot that illustrated the diagnosis ability of a binary classifier system as its discrimination threshold was varied [36]. The ROC curve was created by plotting the true positive rate (TPR) against the false positive rate (FPR) at various threshold settings [36]. The true positive rate was known as sensitivity and the false positive rate was equal to 1-specificity.

To show more performance details of our models and human readers, the confusion matrix was employed to illustrate the prediction results of all three Nugent groups. In the confusion matrix, each row of the matrix represented the instances in a predicted class while each column represented the instances in an actual class (or vice versa) [37]. The accuracy of three groups of classifications was also provided.

### 2.5 Comparison to Human Readers

Three data sets Γ, Λ, Σ were then evaluated by our CNN model and five independent healthcare providers (HCPs), including three technologists and two obstetricians. The five HCPs were from four representative teaching hospitals and the leading private hospitals in China: The First Affiliated Hospital of Xi’an Jiaotong University, The Second Affiliated Hospital of Soochow University, Beijing HarMoniCare women’s and children’s Hospital, Binzhou Medical University Hospital, and Nanjing Medical University Affiliated Nanjing Maternity and Child Health Care Hospital. The diagnostic results were compared using the metrics method.

## 3 Results

### 3.1 Development and Selection of the Best CNN Model for Nugent Scoring

Image data from Beijing Tsinghua Changgung Hospital were used to select the best CNN model for BV diagnosis. Four CNN models were generated and trained by the 23280 training image set, then evaluated on the 5815 test images. For the purpose of screening, we considered altered vaginal flora and BV as positive samples and normal vaginal flora as negative samples. All models had AUCs greater than 0.95, with the 1/4 NugentNet model shown the best performance of AUC = 0.978 (Fig.4(a)). The NugentNet and 1/2 NugentNet had more training variables than the best model. These two models showed overfitting on our data set, and hence gave lower AUCs than the 1/4 NugentNet. The number of training variables of 1/8 NugentNet was only a quarter of 1/4 NugentNet. The AUC of the 1/8 NugentNet was slightly lower (0.004) than the 1/4 NugentNet. Therefore, it was likely underfitting on our data set. The 1/4 NugentNet was chosen for the following study as the best CNN model for the prupose of Nugent scoring and diagnosis of BV.

**Figure 4:**
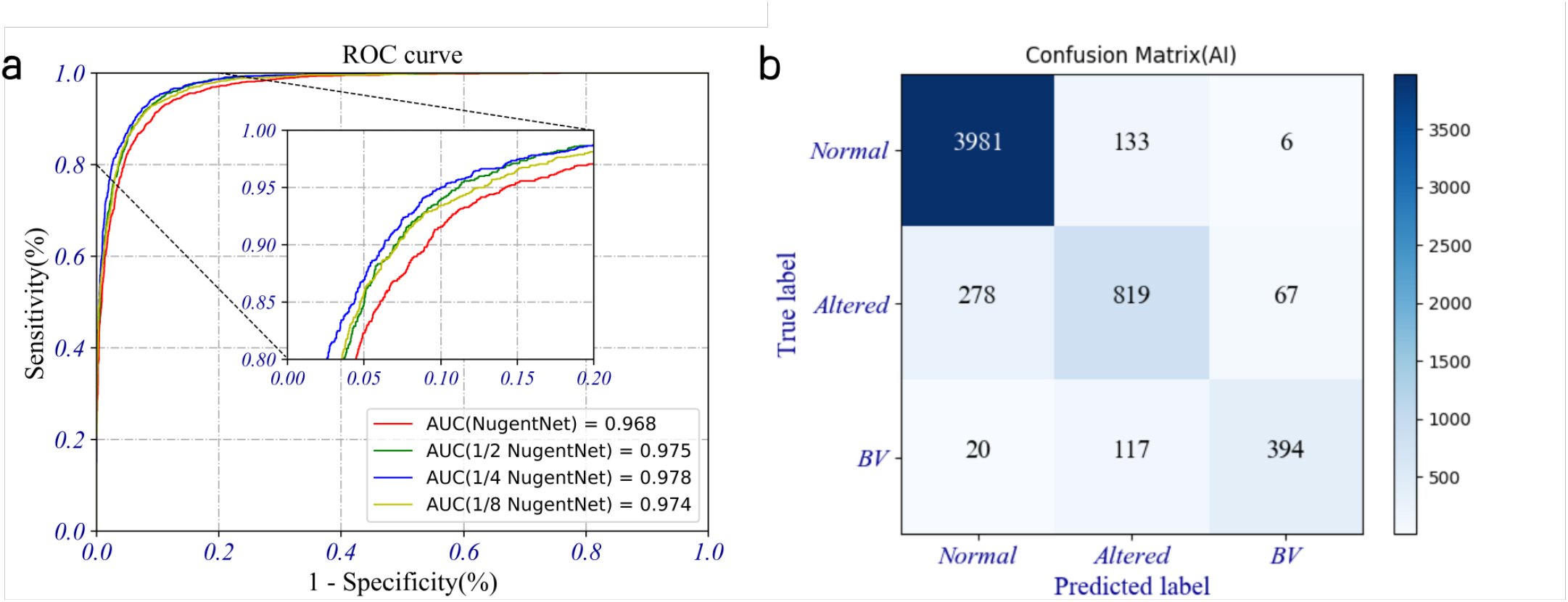
The performance of our models on the test set. a) was the ROC curves of our four models, the 1/4 NugentNet was the best model getting AUC=0.978. b) was the confusion matrix of the best points of the best model.

To better understand the performance of the best model on BV classification, the three classification results (Confusion Matrix) of the best point of AUC curve for the 1/4 NugentNet was plotted against the labeled results of the test set. The best point obtained 89.3% accuracy on the three-category classification, which was 5.6% higher than the microscopists and comparable to the top experts’ performance in China [17]. The results showed only 3.8% (20/531) BV samples were predicted as false negative (normal vaginal flora) and 0.1% (6/4120) normal vaginal flora samples were predicted as false positive (BV).

### 3.2 Performance Comparison Between CNN Model and Human Practitioners

The 1/4 NugentNet model was comparable to Chinese top healthcare providers in classify Nugent scores from microscopic images. To compare our best model and top level healthcare providers, five human readers from 5 representative teaching hospitals and 1/4 NugentNet independently scored test Set Γ, 427 images from Beijing Tsinghua Changgung Hospital. Our model obtained AUC=0.9746, which out performed 4 human readers and comparable to the best human practitioner(Fig.5).

**Figure 5:**
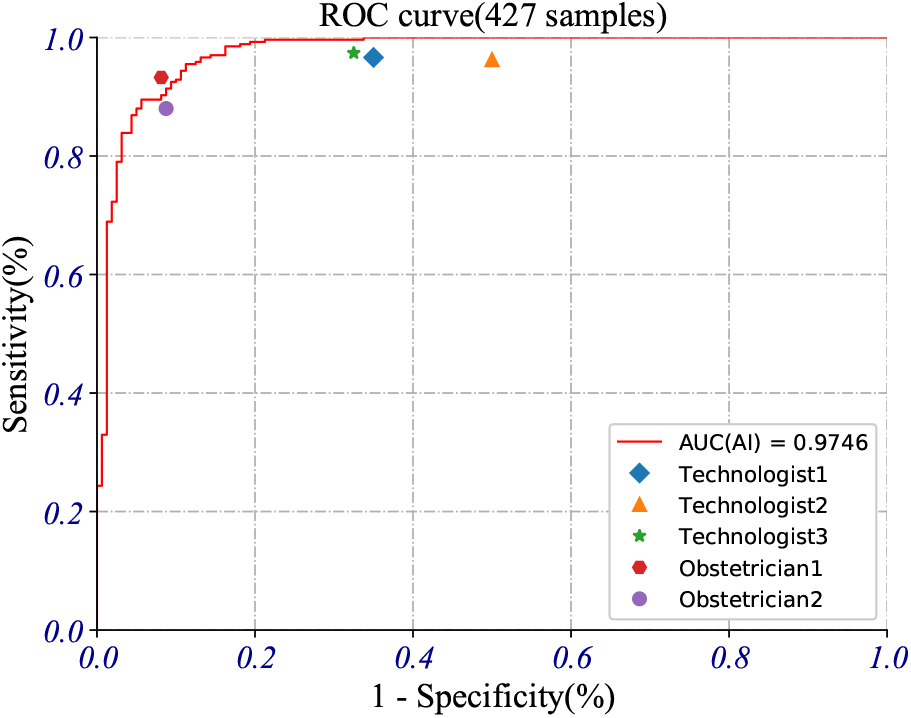
Comparing the results of the best model and five human readers on the independent test set Γ.

The 1/4 NugentNet model had better accuracy in scoring image samples than human practitioners. Because the Nugent scores are categorized in three classifications, the AUC curve does not refect the reality of diagnostic accuracy in clinical settings. The three classification accuracies were calculated for 1/4 NugentNet and 5 human readers. The three-class accuracy of 1/4 NugentNet achieved 80.3%, which was 10.3% higher than the human’s average result and 0.6% higher than the the best human result (Tab. 3). The average performance of all the human readers was 94.3% sensitivity and 73.1% specificity. At the best point of AUC curve, our model had a sensitivity of 91.4% and sensitivity of 91.3%. If we moved the AUC curve of our model at the sensitivity of 94.3% to match the average of human readers, our specificity was 89.4%, a 16.3% higher than the 73.1% average specificity of human readers. Overall, our CNN model achieved high accuracy on Nugent scores from high quality images obtained from Beijing Tsinghua Changgung Hospital.

**Table 3:** The detail information of the results of the best model and five human readers on the independence test set Γ.

|  | Sensitivity | Specificity | Three Classification Accuracy |
| --- | --- | --- | --- |
| Best Points(AI) | 91.4% | 91.3% | 80.3% |
| High Sensitivity(AI) | 97.4% | 83.8% | 80.8% |
| Technologist1 | 96.6% | 65.0% | 66.0% |
| Technologist2 | 96.3% | 50.0% | 62.1% |
| Technologist3 | 97.4% | 67.5% | 67.5% |
| Obstetrician1 | 93.3% | 91.9% | 79.6% |
| Obstetrician2 | 88.0% | 91.3% | 74.7% |
| Average (Technologists) | 96.8% | 60.8% | 65.2% |
| Average (Human Readers) | 94.3% | 73.1% | 70.0% |

### 3.3 Performance of CNN Model on Images from Different Clinical Settings

Since images from different clinical settings differented in size, brightness, and color (Tab.2), the preprocessing procedure was essential to improved the accuracy for our CNN model. Using various preprocessing steps, we had much better performance on data Set Λ and Σ (Tab.4). Data Set Λ had larger actual physical area and brighter samples than the training data set we obtained from Beijing Tsinghua Changgung Hospitals. Adjusting physical area had improved the AUC from 0.8552 to 0.9375. Adjusting brightness had improved the AUC from 0.8552 to 0.8626. Combined the adjustments of brightness and physical area made the performance of our CNN model on Set Λ to an overall AUC of 0.9396 (Tab.4). The test results of Σ showed similar pattern that adjusting of both brightness and physical area had greatly improved the performance, AUC from 0.5137 to 0.9450 (Tab.4). Since images in Set Σ had smaller physical area than the training data, we had to expand the edges of images to match the actual physical area for our model. Of the three common methods tried, our results showed reflect was the best edge expansion method, which had increased the AUC from 0.5137 to 0.7136 (Tab.5). Both adjusting physical area and adjusting brightness greatly improved model performance on the testing data Set Λ and Σ from different clinical settings.

**Table 4:** The performance of the best model using preprocessing on test sets Λ and Σ.

| | Independent Test Set $\Lambda$ | | | | Independent Test Set $\Sigma$ | | | |
| --- | --- | --- | --- | --- | --- | --- | --- | --- |
|  | Original Image | Adjust Physical Area | Adjust Brightness | Adjust Both | Original Image | Adjust Physical Area | Adjust Brightness | Adjust Both |
| AUC | 0.8552 | 0.9375 | 0.8626 | 0.9396 | 0.5137 | 0.7136 | 0.8894 | 0.9450 |

**Table 5:**
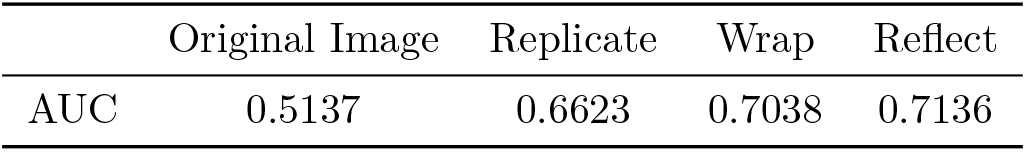
The performance of three edge expansion methods in adjusting physical area step on Σ.

|  | Original Image | Replicate | Wrap | Reflect |
| --- | --- | --- | --- | --- |
| AUC | 0.5137 | 0.6623 | 0.7038 | 0.7136 |

### 3.4 Comparison Between CNN Model and Human on Combined Images

Our model was comparable to human practitioners on Nugent score diagnosis using images from various clinical settings. Data set Γ, Λ, and Σ were combined to get a independent testing data set of 1082 images. When applied the 1/4 NugentNet model on the independent testing data set, it obtained AUC of 0.7007 on original data, 0.7884 after adjustment of physical areas, 0.8917 after adjustment of brightness, and 0.9400 by preprocessing both (Fig. 6). When compared with 5 experts who diagnosed the Nugent scores on the same image set, our model out performed three of them judged by sensitivity and specificity (Fig. 6). The average performance of all the technologists was 96.5% sensitivity and 62.2% specificities. When setting the sensitivity equal, our model had a specificity of 71%, 8.8% higher than the technologists’ average results. When specificity was setted equal, the sensitivity of our model was 2.1% higher than technologists’ average. Using the three-class accuracy as better read out of the Nugent scoring performance, our 1/4 NugentNet model achieved a total accuracy of 75.1%, which is again better than three human experts and 2.1% higher than human average (Tab. 6). The results showed that our model had great performance by applying preprocessing steps to different images.

**Table 6:** The detail information of the results of the best model and five human readers on the total independent test sets with 1082 samples.

|  | Sensitivity | Specificity | Three Classification Accuracy |
| --- | --- | --- | --- |
| Best Points(AI) | 89.0% | 85.0% | 75.1% |
| High Sensitivity(AI) | 97.3% | 69.3% | 71.1% |
| Technologist1 | 97.0% | 60.0% | 67.8% |
| Technologist2 | 95.3% | 60.9% | 67.5% |
| Technologist3 | 97.3% | 65.6% | 70.3% |
| Obstetrician1 | 94.4% | 93.9% | 80.9% |
| Obstetrician2 | 90.4% | 92.7% | 78.7% |
| Average (Technologists) | 96.5% | 62.2% | 68.5% |
| Average (Human Readers) | 94.9% | 74.6% | 73.0% |

**Figure 6:**
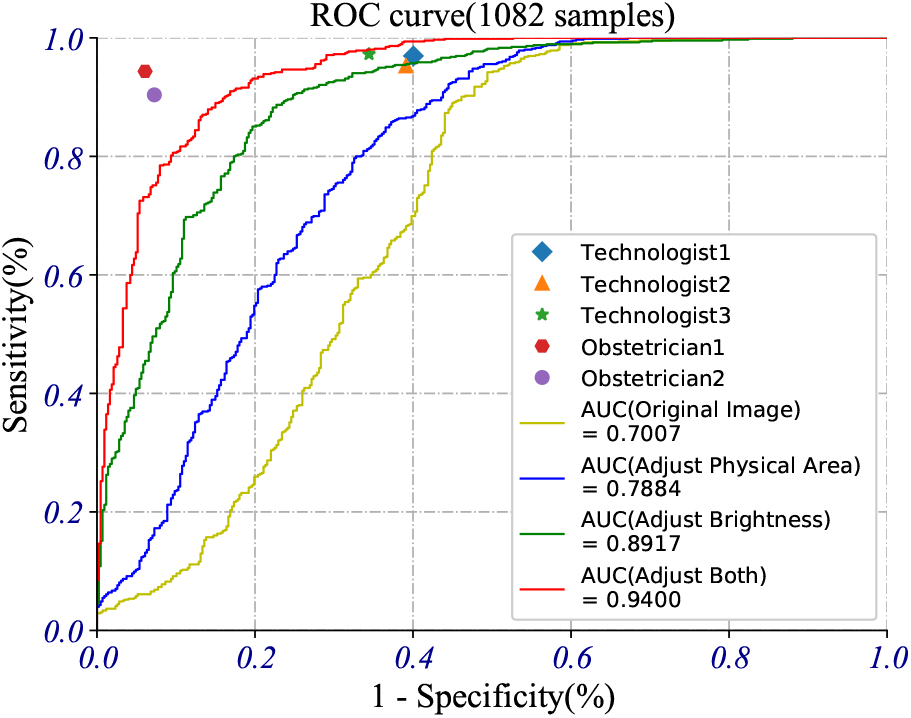
Comparing the results of the best model and five human readers on the total independent test sets with 1082 samples.

## 4 Discussion

In the past years, CNN methods had shown lots of examples of successful applications in medical image processing. Typical examples were: automatically classification 26 common skin conditions [39]; using pixels and disease labels as inputs to classify skin lesions [40]; identified prostate cancer in biopsy specimens and detected breast cancer metastasis in sentinel lymph nodes [41]; automated classification of Gram staining of blood samples [33]; automatically diagnosis of H. pylori infection [32]; predicting cardiovascular risk factors from extracted retinal fundus images [31]. Therefore, the CNN methods was suitable for medical image processing. As a type of medical image, the microscopic image contained lots of local details and global information. The local details included various types of bacteria. The global information included the distribution density and ratio of various bacteria, etc. In the diagnostic process, all the information should be considered for an accurate result. The local details could be extracted by the first few layers of the CNN model, while the global information could be extracted by the last few layers. The last fully connected layer could use all the information extracted from the convolution layer in front to obtain the Nugent score. The CNN model could accurate extract local and global information for diagnosis [27,28,38]. Therefore, the CNN model was very suitable for Nugent score automatic diagnosis.

The 1/4 NugentNet CNN model showed better performance to process microscopic images for BV diagnosis using Nugent score than traditional image processing method. Traditional automatic image processing methods required three difficult diagnostic steps to get the diagnosis results [17]. The first step involved the segmentation of the bacterial area, which required a series of artificially designed algorithms to extract the foreground. In the second step, the overlap clumps would be split from the bacterial area, from which morphotypes of individual bacterium were obtained. In the third step, the features of bacterial morphotypes were extracted and classifed using traditional machine learning methods. In our CNN model, the features of the microscope images could be automatically extracted with diagnosis made spontaneously, which avoided complex diagnostic steps in the traditional methods. The sensitivity, specificity and three-class accuracy of traditional method were only 58.3%, 87.1% and 79.1% [17]. Our 1/4 NugentNet model had better performance in all the three diagnostic performance indicators: a 24.1% increase for sensitivity, 9.5% increase for specificity, 10.2% increase for three-class accuracy. Furthermore, our model could simultaneously adjust the sensitivity and specificity by adjusting the predicting probability threshold. The diagnostic performance could not be further improved with the same traditional automatic diagnostic methods, but could be further improved in our model with more training data.

The diagnosis accuracy of 1/4 NugentNet CNN was comparable to human experts with advantages in consistency and flexibility. The test data set of 1082 images were judged by top experts, with each result agreed by at least two of them. When tested independently by our CNN model and 5 human experts, results of 1/4 NugentNet on the test data set had an average three-class accuracy 75.1% which is 2.1% higher than the average of experts. Although the overall accuracy seems comparable, human readers showed considerable variability of sensitivity from 90.4% to 97.3%, and big variation in specificity from 60.0% to 93.9% (Tab. 6). When we considered the most accurate point of our CNN model, the specificity and sensitivity were 85.0% and 89.0% respectively. Aplication of CNN model in clinical settings could minimize the potential diagnostic variations brought by different practitioners. Adjusting on the AUC curve, we could also find diagnostic point with specific sensitivity for the purpose of clinical practice. For screening purpose, we could increase the sensitivity to identify positive patients as many as possible. For confirmative testing, we could increase the specificity to minimize false diagnosis. The CNN model could be implemented with different capacity in clinical settings.

Our model showed great performance and was overperformed technologists when the samples was standardized by the preprocessing. Standardizing the actual physical area and the brightness of the samples made our model perform very well on variable samples from different settings. We further studied the impact of the clarity of images and the number of training set on the accuracy of our models. The results showed the image sharpening method did not improved the model’s performance on the independent test sets, and the performance decreased when the samples were blurred. The clarity of the samples was good enough for our models. To investigate the performance of our best model with different training sample sizes, we trained the best model with five different sample sizes including 5000 images, 10000 images, 15000 images, 20000 images, and 23000 images. All the AUCs were greater than 0.95 (Fig.7). As we expected, a larger training set could produce a model with better performance. When the training set had more than 15000 samples, the performance of the model would improved only marginally as the number of samples increased.

**Figure 7:**
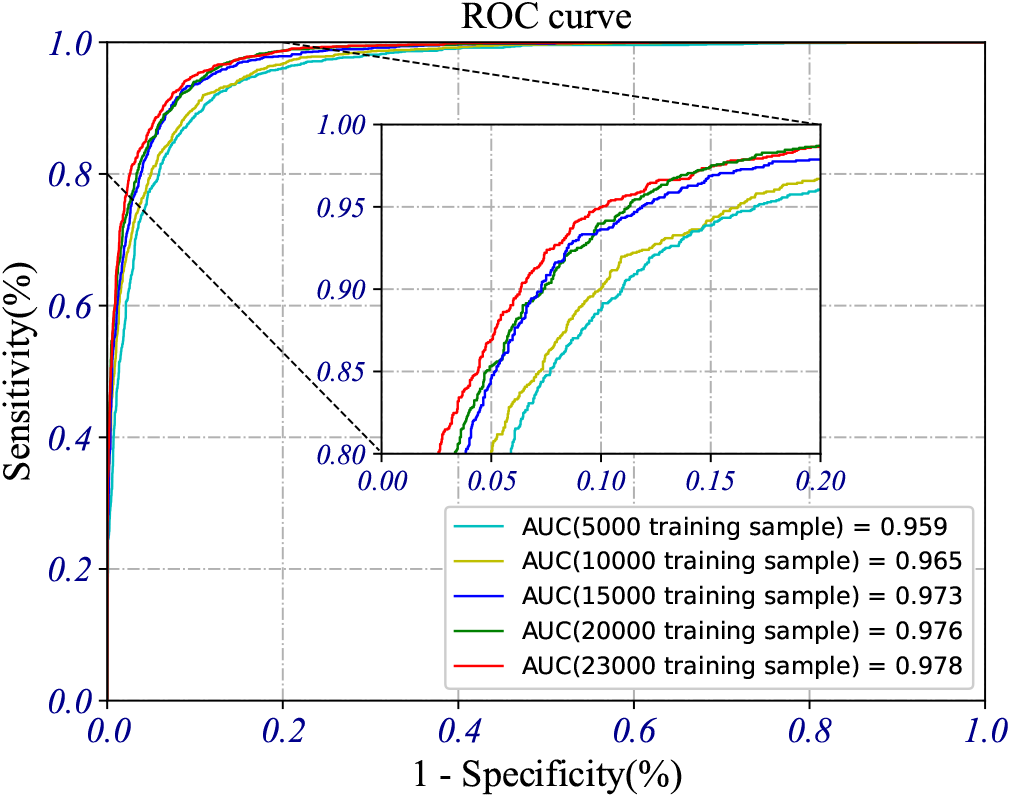
The performance of the best model with different training sample sizes: 5000 images, 10000 images, 15000images, 20000 images, and 23000 images.

The CNN model can increase the speed of BV diagnosis in clinical settings. We used a NVIDIA GeForce GTX 1080Ti GPU for training and inference. The best model was faster than the NugentNet and 1/2 NugentNet in both training process and inference process. In the training process, the 1/4 NugentNet model could be obtained within 10000 iterations and could be completed in 2.4 hours by one GPU. But it needed months to years to train an proficient technologist. In the inference process, it took our model 2.4 seconds to diagnose 100 images, while the traditional automatic diagnostic methods needed 30 seconds to obtain the diagnosis result for a single microscope image [17]. By using the same hardware, our model could diagnose 5 microscopic images per second. The inference speed of our model was more than 150 times faster than traditional automatic diagnostic methods. The diagnostic efficiency of our model was much higher than traditional automatic diagnostic methods.

In conclusion, our study first proved deep learning techniques to diagnose bacterial vaginosis based on microscopic images. we constructed the convolutional neural network models for automatic BV diagnosis. For image-level BV diagnosis, our models had better performance in terms of 3-class accuracy (89.3%) than experts and traditional automated diagnostic methods. With the help of automatic scanning microscope, manual nugent scoring can be completely replaced by our model and the problem of clinical manual Nugent scoring can be completely solved. In addition, lots of gynecological lower genital tract infections including aerobic vaginitis (AV), vulvovaginal candidiasis (VVC) and trichomonas vaginitis (TV) were diagnosed by the same microscope images. Our model could be further modified to diagnose all three infections. Furthermore, our model could be used for diagnosis of other microscope images for infection diagnosis in different clinical settings such as sepsis. It could be developed into an automatic diagnostic device for more precise, more efficient and more stable than manual diagnosis and would standardize diagnostic process.

## Data sharing statement

The data that support the findings of this study are available from the corresponding author upon reasonable request.

## Author contributions

Z. Wang, L. Zhang, Z. Liu, Q. Liao and W. Xu proposed the research, Z. Liu, R. An, P. Li, L. Geng, Q. Qiao, W. Zhu and Q. Liao led the multicenter study, Y. Wang, Z. Wang, and L. Qi collected data, Y. Wang, H. Bai, and M. Zhao performed the data annotation, Z. Wang, Z. Yang, Y. Cao, M. Li and W. Wu wrote the deep learning code and performed the experiment, J. Li, N. Li, C. Rui, C. Fan, X. Liu, Y. Si, L. Qi and A. Feng evaluated the algorithm, Z. Wang, L. Zhang and Q. Zhang wrote the manuscript, M. Wang, W. Mo, Q. Liao and W. Xu reviewed the manuscript.

## Acknowledgment

Thanks to Suzhou Turing Microbial Technologies Co., Ltd and Beijing Turing Microbial Technologies Co., Ltd for technical support.

